# Cell-to-cell variations in subcellular localisation of V2R are independent of its expression level

**DOI:** 10.1101/2021.08.09.455709

**Authors:** Eline J. Koers, Bradley A. Morgan, Franziska M. Heydenreich, Bianca Plouffe, Iain B. Styles, Dmitry B. Veprintsev

## Abstract

G protein coupled receptors (GPCRs) translate the actions of hormones into intracellular signalling events. Mutations in GPCRs can prevent their correct expression and trafficking to the cell surface and cause disease. We use single cell measurements in HEK293 cells to show that the balance between endoplasmic reticulum (ER) and cell surface localisation of the Vasopressin 2 receptor (V2R) varies significantly from cell to cell. We find that mutations in the V2R affect the proportion of cells able to send this GPCR to the cell surface but do not prevent all cells in the population from correctly trafficking the mutant receptors. These findings reveal that the ability of cells to correctly traffic V2R to the cell surface depends not only on the expressed V2R mutant but also on the individual cell environment.

**Significance statement:** Missense mutations in the Vasopressin 2 Receptor (V2R) cause Nephrogenic Diabetes Insipidus. Some of these mutations prevent correct expression and trafficking of V2R to the cell surface resulting in a loss-of-function. We show -using single cell measurements-that the balance between endoplasmic reticulum and cell surface localisation of the V2R varies significantly from cell to cell, independent from its expression level. Mutations affect the proportion of cells able to send V2R to the cell surface but do not prevent all cells in the population from correctly trafficking the mutant receptors. Hence, the ability of cells to correctly traffic V2R to the cell surface depends not only on expressed V2R mutant but also on the cell environment.

## Introduction

G protein coupled receptors (GPCRs) coordinate functions of multicellular organisms by sensing hormones, neurotransmitters or cytokines that circulate around the body and inducing intracellular responses. Missense mutations in these receptors often cause loss-of-function and are linked to 55 different monogenic diseases (Schöneberg, Liebscher, & Insel, 2020).

Missense mutations in GPCRs can lead to a change or loss of ligand binding and intracellular signalling (i.e. function). Alternatively, missense mutations disrupt receptor biogenesis. GPCRs are membrane proteins that canonically initiate their signalling at the cell surface. Alterations in expression levels and copy number of receptors at the cell surface can cause dysfunction in their intracellular signalling responses.

The Vasopressin 2 Receptor (V2R) is mainly known for its function in the kidney where it promotes translocation of the aquaporin 2 channel to the apical plasma membrane of the epithelial cells of the collecting duct, resulting in water reabsorption (Wilson, Miranda, & Knepper, 2013). Many mutations in the Vasopressin 2 Receptor (V2R) are associated with NDI (nephrogenic diabetes insipidus) and cause a loss-of-function phenotype, preventing the kidneys’ normal response to vasopressin. Several deletions and mutations have been reported to cause accumulation of the receptor in the endoplasmic reticulum (ER) (Robben, Knoers, & Deen, 2005). Addition of a pharmacological chaperone, for example the antagonist SR121463, can restore cellsurface expression for some mutants by stabilising trafficking-competent receptor conformations (Morello et al., 2000). Accumulation in the ER is thought be caused by misfolding, maturation defects and increased interaction with ER-resident proteins such as calnexin. (Morello et al., 2001). Protein degradation pathways also may play an important role in ability of GPCRs to reach cell surface, exemplified by the proteosome inhibitor bortezomib being able to rescue cell-surface expression of ER-retained mutants of GPCRs in a universal matter (Morfa et al., 2018). Many potential effectors that are involved in GPCR folding and quality control may affect trafficking of GPCRs to the cell surface (Achour, Labbé-Jullié, Scott, & Marullo, 2008) and a dependence on the cell type with respect to the subcellular localisation of a GPCR has been reported (Blagotinšek Cokan et al., 2020).

Given the stochastic nature of gene expression in individual cells, including proteins involved in GPCR folding, trafficking and degradation, it is important to ask what produces a more dominant effect: the mutations in GPCRs that trigger ER retention or is it the cellular environment that determines the GPCR fate?

We hypothesise that the interplay of a specific mutation and the cellular environment is a strong factor in determining receptor fate and biogenesis efficiency. To test this, we systematically probed a large panel of mutants for expression and localisation in the same cell type. We measured individual cell outcomes to probe variations in a specific cell type population.

We chose to study the expression and subcellular localisation of a number of variants of vasopressin 2 receptor (V2R) identified by alanine scanning mutagenesis as having reduced cell surface expression (Heydenreich, 2016), complemented by a panel of NDI-causing mutants.

We show that the majority of the 28 alanine mutations and 6 NDI-causing mutations we studied lead to the retention of V2R in the ER, preventing its correct trafficking to the cell surface suggesting that the mutations likely affect biogenesis or folding, which is consistent with prior observations. Overall, at the cell population level, we observe that mutations affect subcellular localisation and expression level. However, further analysis of subcellular localisation in individual cells suggest that within each population, individual cells show highly varying levels of ER localisation of V2R which is not correlated with total expression levels. While the molecular basis for existence of these distinct “folding and trafficking” states of cells is not clear, it does suggest that there are some common factors that could be controlled for therapeutic purposes. Given the strong structural conservation among GPCRs, our findings are likely to be applicable to many other GPCRs where mutations cause diseases and open novel questions in the field of GPCR biogenesis.

## Results

### Most tested V2R mutants are more localised in the ER than V2R Wild Type

We examined whether common NDI-causing mutations (fig 1A) and synthetic variants (i.e. alanine scanning mutants) with low surface expression levels (<25% of WT) (Fig 1B) (Heydenreich, 2016) have early biogenesis defects e.g. incorrect topology, misfolding or structural instability, and are held in the ER as a result.

**Figure 1.**
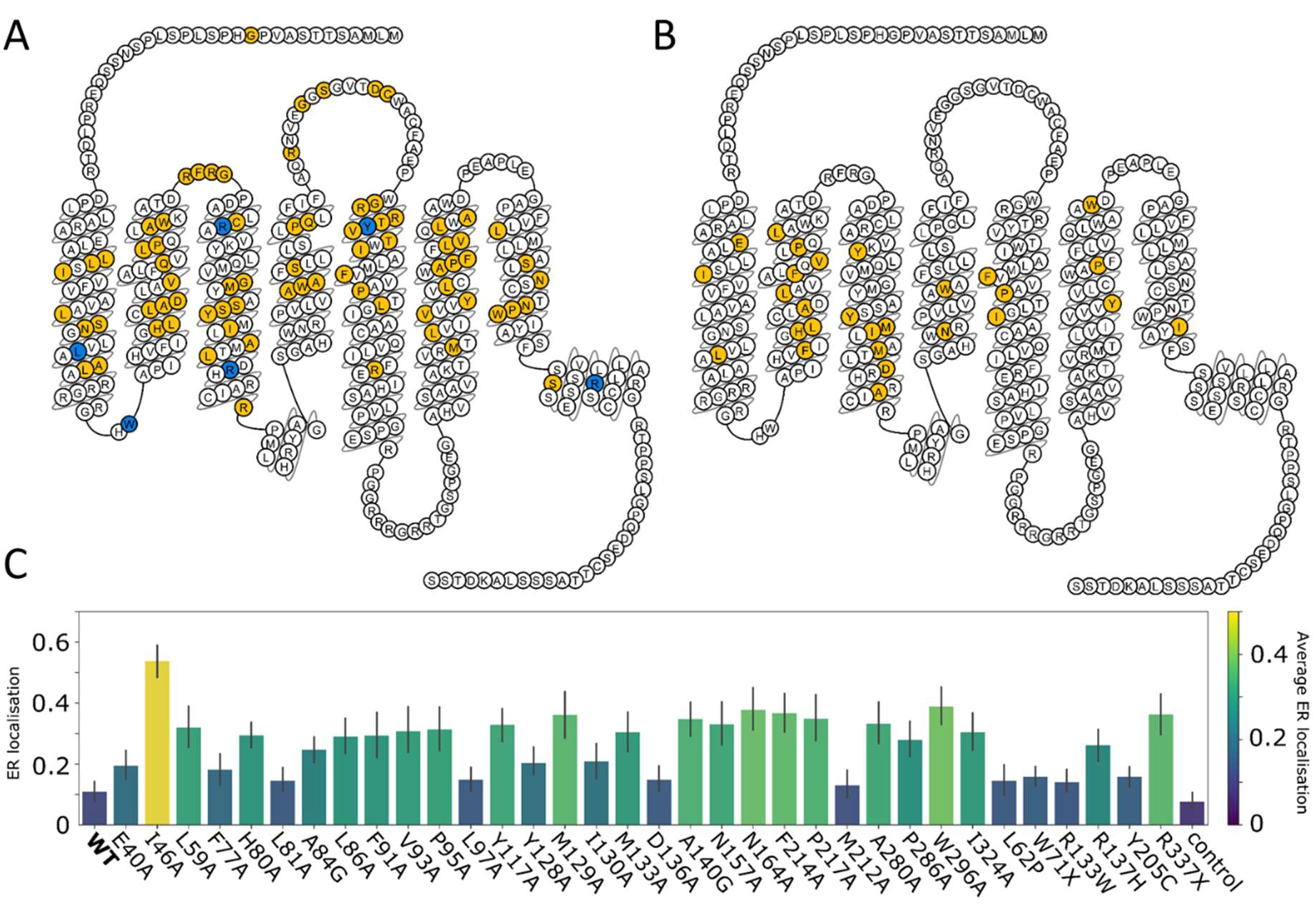
Structural position of NDI-causing and cell-surface expression reducing mutations in V2R, and their ER retention. A) NDI-causing missense mutations (Spanakis, Milord, & Gragnoli, 2008) shown in yellow and common NDI-causing mutations in blue B) Alanine scanning mutants with low (<25% of WT) cell surface expression in yellow (Heydenreich, 2016). C) Subcellular localisation of low expressing mutants (averaged across all images) varies between cell surface and ER. Depicted are average numbers and 95% confidence interval.

We used confocal microscopy combined with immunostaining to test this. We determined the ER localisation by means of calculating the Manders’ overlap coefficient over two experiments and 16 images per mutant. Manders’ overlap coefficient reports on coincidence detection of two coloured proteins in the same pixel of an image. For example, if the ER marker is detected in 100 pixels of the image, and the V2R is detected in all of these 100 pixels, the Manders’ score would be 1. If V2R is detected in other pixels and no coincidence detected, it will be 0. Most of the alanine mutants (24/28 see table S1) show higher presence in the ER than wild type (WT) (p<0.05), pointing to early biogenesis defects as a cause for lower cell surface expression and/or dysfunction (Fig. 1C).

We investigated whether the studied mutations are predominantly found in the ligand binding pocket or G-protein binding sites, or outside of these regions. We found that 27% of NDI-causing missense mutations and 14% of the low surface expression alanine mutations were in functional sites. For both sets, there is no significant (p > 0.05) difference with a random distribution of mutations.

### Single cell analysis of subcellular localisation shows different rates of ER retention for cells under similar conditions

We observed a large spread of measured ER retention values when analysing images as a whole, as well as visually noticeable differences in subcellular localisation of V2R in individual cells (Fig 2A,B). Hence, we decided to analyse the images further on a single-cell level. We set up an analysis pipeline using CellProfiler (details in Methods section) (Misteli et al., 2018) and determined ER localisation and overall fluorescence intensity per cell. We found that the averages from all images and per cell analysis were similar to the whole image analysis, while the *per cell* analysis shows a wider spread of ER localisation of cells grown under the same conditions expressing the same V2R mutant (Fig 2C).

**Figure 2.**
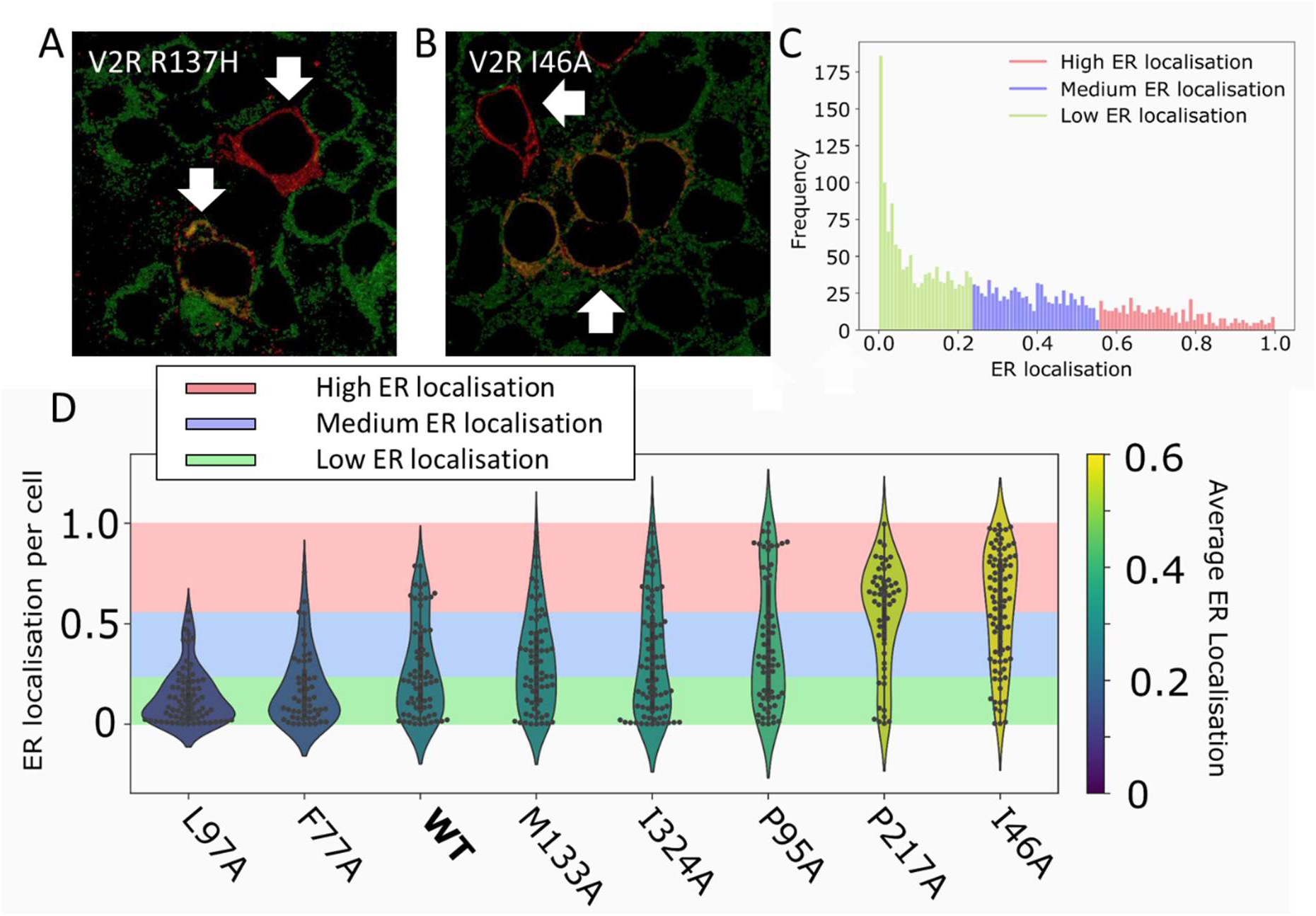
Single cell analysis of ER localisation. A,B) Examples of fluorescence images of cells transfected with two different V2R mutants. ER (green) and V2R (red) showing differences in ER localisation. C) Subclasses of all cells based on ER localisation D) ER localisation (MOC) of a selection of alanine mutants per individual cell. The cells are divided in three categories: low, medium and high ER localisation based on k-means clustering. ER localisation data of the other alanine mutations can be found in figure S1.

In order to examine the ER localisation distribution further, we used k-means analysis to divide the cells in three categories, high, medium and low ER localisation (Fig 2B,C).

High protein expression levels can cause ER stress and saturate the folding and trafficking machinery, resulting in higher ER retention of V2R.

When we examined the average fluorescence intensity, we indeed observed that some mutants were expressed in lower amounts than WT, while others had higher expression (fig 3B). We further explored whether there was any correlation between the intensity – reflecting the protein expression level – and high ER localisation. The possibilities include high concentration of the receptor overloading the ER trafficking capability. Alternatively, low levels of protein may be due to efficient degradation in the ER. However, we found no correlation between V2R mutant expression and ER localisation on the individual cell level (fig S3).This means that differences are caused by other – potentially subtle – differences between cells.

**Figure 3.**
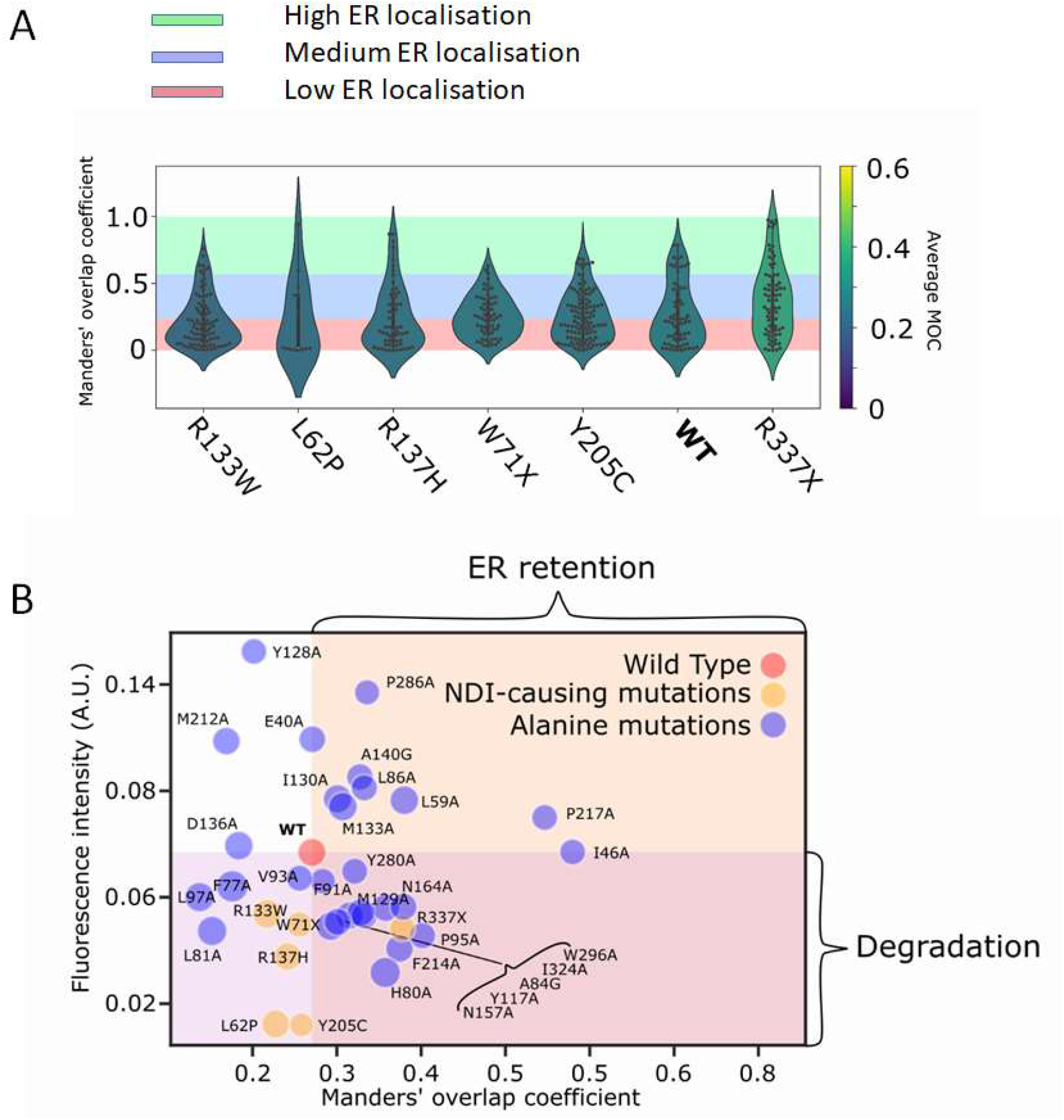
Single cell ER localisation of V2R patient mutations and average intensity and ER localisation of all cells. A) Per cell ER localisation (MOC) of NDI-causing patient mutations compared to V2R WT. All V2R patient mutations tested except V2R R337X show lower ER localisation than V2R WT, B) Average V2R expression (fluorescence intensity) and ER localisation (MOC) of all studied mutants including V2R WT (red), Alanine mutants (blue) and NDI-causing mutations (yellow). The pink shaded area denotes lower fluorescence intensity than V2R WT which signifies lower expression levels or degradation. The orange shaded area marks higher ER localisation than V2R WT which signifies ER retention.

Alternatively, we hypothesized that *per cell* ER localisation may be influenced by the cell-cycle phase. We examined this by measuring the intensity of the nuclear stain DAPI that was shown to be a very predictive parameter to identify cell-cycle phase (Ferro et al., 2017). We found that >95% of the cells are in same cell-cycle phase, i.e. G0 (fig S4). V2R WT and V2R E40A show a subset of cells with a higher concentration of DNA in the nucleus indicating that some cells are in S, G2 or M. Hence, cell cycle phase cannot explain the ER localisation differences observed.

### V2R patient mutations show moderate presence in ER but lower overall expression on a *per cell* basis

We tested whether six common NDI-causing mutations (Spanakis et al., 2008) showed an increase in ER retention compared to WT V2R. We found that, apart from a small increase for R337X, none of the other mutants showed higher ER retention (fig 3A). We did find a weaker fluorescence intensity for all patient mutations (fig 3B, orange), which could indicate that these mutants are degraded at a higher rate.

## Discussion

Here, we show the potential of mutations in the V2R to affect the subcellular distribution of protein and cause ER retention. Mutations also affect the level of protein expression.

However, we also observed that the mutation is not the only factor influencing the ability of the receptor molecule to be correctly trafficked by the cells. Individual cells expressing the V2R or its mutants can be divided into subclasses. The first subclass of the cells in the population can express proteins normally and traffic receptors to the cell surface. The second subclass of cells retains receptors in the ER, and a third subclass shows the receptor distributed between the ER and the cell surface. We observed this for the WT protein as well as for mutants, however the relative proportions of cells belonging to these sub-classes varied depending on the mutant expressed. We also observed that not only the subcellular localisation but also the level of expression of individual V2R mutants also varied on a per cell basis, as well as per population.

Where could this variability of protein level and subcellular localisation originate from? A number of mRNA molecules and correspondingly, polypeptide chain production per cell can vary from cell to cell, resulting in variations in protein abundance. A common concept of “ER jamming” suggests that high levels of protein expression may lead to saturation of the ER folding and trafficking machinery, and lead to increased ER retention. However, we found no correlation between protein expression level and ER retention at a single cell level, suggesting that at least under our experimental conditions we have not overloaded the ER. While cell cycle phase could also influence protein expression, in our experiments the majority of the cells were in the same G0 phase.

This leaves options of as yet unidentified factors affecting the efficiency of processes of membrane insertion and folding, or quality control (QC), degradation machinery, or trafficking machinery. The protein abundance of these factors may also vary cell-to-cell, resulting in these different “folding and trafficking” sub-classes. Possible analogy with the functioning of the much better understood soluble folding processes can help us to generate future hypotheses to investigate. In the case of the Hsp90/70 chaperone system, FANCA proteins with severe and function-disruptive mutations are more likely to be associated with Hsp70 while more mild function-permissive mutants are associated with Hsp90 (Karras et al., 2017). Whether a similar functional division is present in membrane proteins remains to be studied.

The NDI-causing V2R mutants we probed did not show more ER localisation compared to the WT V2R. This is in contrast with previous reports (Morello et al., 2000). However, the construct used here contains a soluble N-terminal domain -a SNAP tag-which could influence the insertion pathway of GPCRs depending on V2R’s dependency of insertion by EMC (Chitwood, Juszkiewicz, Guna, Shao, & Hegde, 2018).

One of the direct consequences of our observations is that the otherwise identical cells in tissues would have different levels of cell surface receptors, and will respond differently to changes in hormone and drug concentrations. The importance of this for whole tissue pharmacological response is not clear but should be considered in future studies. The other exciting aspect of this work is the finding that even rather disruptive mutants can be produced and trafficked to the cell surface suggesting that the process of biogenesis can be manipulated by targeting proteins –e.g. proteostasis modulators-involved in this process to increase overall production of the mutant receptor, resulting in (partial) restoration of its function. However, the next challenge is to identify the key players in V2R biogenesis that affect the efficiency of this process.

## Acknowledgements

We thank Jeremy Pike, Clare Harwood, David Sykes and Michel Bouvier for valuable advice and critical discussions and Tim Self, Seema Bagia and Robert Markus from the SLIM Imaging facility at the University of Nottingham for their support. BAM is funded by the MRC IMPACT doctoral programme. EK was funded by the IBSA foundation for scientific research. BP was supported by a Postdoctoral Fellowship from Diabetes Canada and an Early-Career Small Grant for Basic Scientists (10/0006000).

## Materials and Methods

### Distribution of mutations in and outside functional sites

The ligand and G protein binding sites were defined using structural information derived from the 7HK0 cryo-EM structure of V2R in complex with vasopressin and G-protein. (Wang et al., 2021) Two-sided Welch’s t-test was done to test whether the distribution of NDI-causing mutations in and outside functional significantly differed from chance distribution. The same was done for the alanine scanning mutations with low surface expression.

### Cloning and selection of V2R mutants

All mutants were cloned into a pcDNA4/TO vector containing an N-terminal Twin-Strep and SNAP tag (Heydenreich et al., 2017). Alanine scanning mutant plasmids were made and obtained from (Heydenreich, 2016) as well as cell surface expression data to select alanine mutants with <25% cell surface expression compared to V2R WT. NDI mutations were cloned using a two-fragment approach (Heydenreich et al., 2017). Primers were designed using the PCRdesign software (Sun et al., 2013). The selected mutants are listed in Table S2.

### V2R mutant expression and immunolabeling

HEK293T cells were grown in 6-well plates with 22 mm square no. 1.5 coverslips coated with poly-D-lysine hydrobromide (Sigma-Aldrich, #P6407). Cells in each well were transiently transfected using 100 ng V2R and 900 ng sheared salmon sperm DNA (Invitrogen) and 1:3 25 kDa linear Polyethylenimine (Polysciences, #23966). The cells were grown for 20-28h. The cells were fixed using 3% paraformaldehyde, incubated for 20 min, and washed 2x with PBS. Coverslips were then elevated from 6-well plates and cells permeabilised with 0.5% IGEPAL in PBS for 5 min at 4°C, washed 2x 5 min with PBS, then blocked with 3% BSA-1% glycine in PBS, incubated for 30 min, washed 2x 5 min with PBS, then blocked with 10% Goat serum (Abcam, #ab7481), incubated 30 min, then added polyclonal anti-SNAP antibody (Thermo-Fisher,# CAB4255) 1:100 and Anti-P4HB antibody [RL90] (abcam, #ab7481) 1:500 in 10% Goat serum and incubated overnight at 4 C. Washed coverslip 3x in PBS and incubated cover slips in goat-anti-rabbit-AF-647 (Abcam, #ab150083) 1:500, goat-anti-mouse-AF-488 (Abcam, #ab150117) 1:500, DAPI stain 5 mg/mL 1:1000 in 10% Goat serum in the dark for 1h. Washed coverslips 3×5min with PBS and mounted coverslips with Vectashield antifade mounting medium (Vectorlabs, #H-1000).

### Microscopy

Images were recorded using a Zeiss LSM 710 laser scanning confocal microscope fitted with a Zeiss Plan-Apochromat 63x/1.40 NA oil immersion objective. A Diode 405 nm, Helium Neon 633 nm laser and Argon laser at 488 nm were used to excite DAPI, Alexafluor 647 and Alexafluor 488 fluorophores respectively and emission was collected using a 488/561/633 multi beam splitting filter. Images were taken at 512 × 512 pixels per frame and a slice of 53.36 um for confocal images. Laser power and gain were kept constant between experiments.

### Whole image analysis

Whole images were analysed using Fiji (Schindelin et al., 2012) and a batch processing script using 1*σ* Gaussian blur, Moments automatic thresholding algorithm and the Coloc 2 plugin to calculate the Manders’ overlap coefficient per image.

### Single cell image analysis

Single cell image analysis was done using a custom analysis pipeline in CellProfiler 3.1.9 (Misteli et al., 2018). First, nuclei were detected based on DAPI fluorescence images, then the ER and V2R fluorescence images were automatically thresholded using the Otsu algorithm and assigned to a nucleus using a Watershed - Image algorithm. Then the nucleus area was subtracted, and co-localisation was measured by calculating the Manders’ overlap coefficient. In addition, total intensity and area of each cell were calculated. Then, individual entries were filtered for entries with Manders’ overlap coefficient of 0 or 1 and area smaller than 4000 pixels.

### Clustering of cell population based on ER localisation

The *single cell* Manders’ overlap scores were standard scaled clustered using k-means from the Scikit-learn library (Pedregosa et al., 2011). The number of clusters was chosen based on the silhouette and elbow method. Breaks were checked using Jenkspy (https://github.com/mthh/jenkspy).

## Supplementary information

**Table S1.** Table accompanies data presented in fig 1C. Two-sided t-test with unequal variances of ER localisation between WT and each mutant

| mutant | p-value | significant p>0.05? |
| --- | --- | --- |
| E40A | 6.0E-03 | Yes |
| I46A | 5.0E-19 | Yes |
| L59A | 2.5E-06 | Yes |
| F77A | 2.1E-02 | Yes |
| H80A | 3.9E-09 | Yes |
| L81A | 1.8E-01 | No |
| A84G | 1.9E-05 | Yes |
| L86A | 1.9E-06 | Yes |
| F91A | 1.5E-04 | Yes |
| V93A | 4.8E-05 | Yes |
| P95A | 6.2E-06 | Yes |
| L97A | 1.4E-01 | No |
| Y117A | 5.5E-08 | Yes |
| Y128A | 1.4E-03 | Yes |
| M129A | 1.4E-06 | Yes |
| I130A | 6.3E-03 | Yes |
| M133A | 5.2E-06 | Yes |
| D136A | 1.6E-01 | No |
| A140G | 8.0E-09 | Yes |
| N157A | 1.8E-06 | Yes |
| W164A | 2.4E-08 | Yes |
| F214A | 9.5E-09 | Yes |
| P217A | 2.0E-06 | Yes |
| M212A | 4.6E-01 | No |
| Y280A | 4.4E-07 | Yes |
| P286A | 9.4E-06 | Yes |
| W296A | 3.8E-10 | Yes |
| I324A | 1.9E-06 | Yes |
| L62P | 2.7E-01 | No |
| W71X | 4.2E-02 | Yes |
| R113W | 2.3E-01 | No |
| R137H | 1.4E-05 | Yes |
| Y205C | 4.9E-02 | Yes |
| R337X | 4.2E-08 | Yes |
| control | 1.5E-01 | No |

**Table S2.** List of the 28 Alanine scanning mutants and 6 missense and nonsense patient mutations studied, Note: X (e.g. in W71X) stands for termination.

| Alanine scanning mutants | Functional site |
| --- | --- |
| E40A | Yes |
| I46A | No |
| L59A | No |
| F77A | No |
| H80A | No |
| L81A | No |
| A84G | No |
| L86A | No |
| F91A | No |
| V93A | Yes |
| P95A | No |
| L97A | No |
| Y117A | No |
| Y128A | No |
| M129A | No |
| I130A | No |
| M133A | No |
| D136A | Yes |
| A140G | Yes |
| N157A | No |
| W164A | No |
| F214A | No |
| P217A | No |
| I221A | No |
| Y280A | No |
| P286A | No |
| W296A | No |
| I324A | No |
| NDI-causing mutations |  |
| L62P | No |
| W71X | - |
| R113W | No |
| R137H | Yes |
| Y205C | Yes |
| R337X | - |

**Figure S1:**
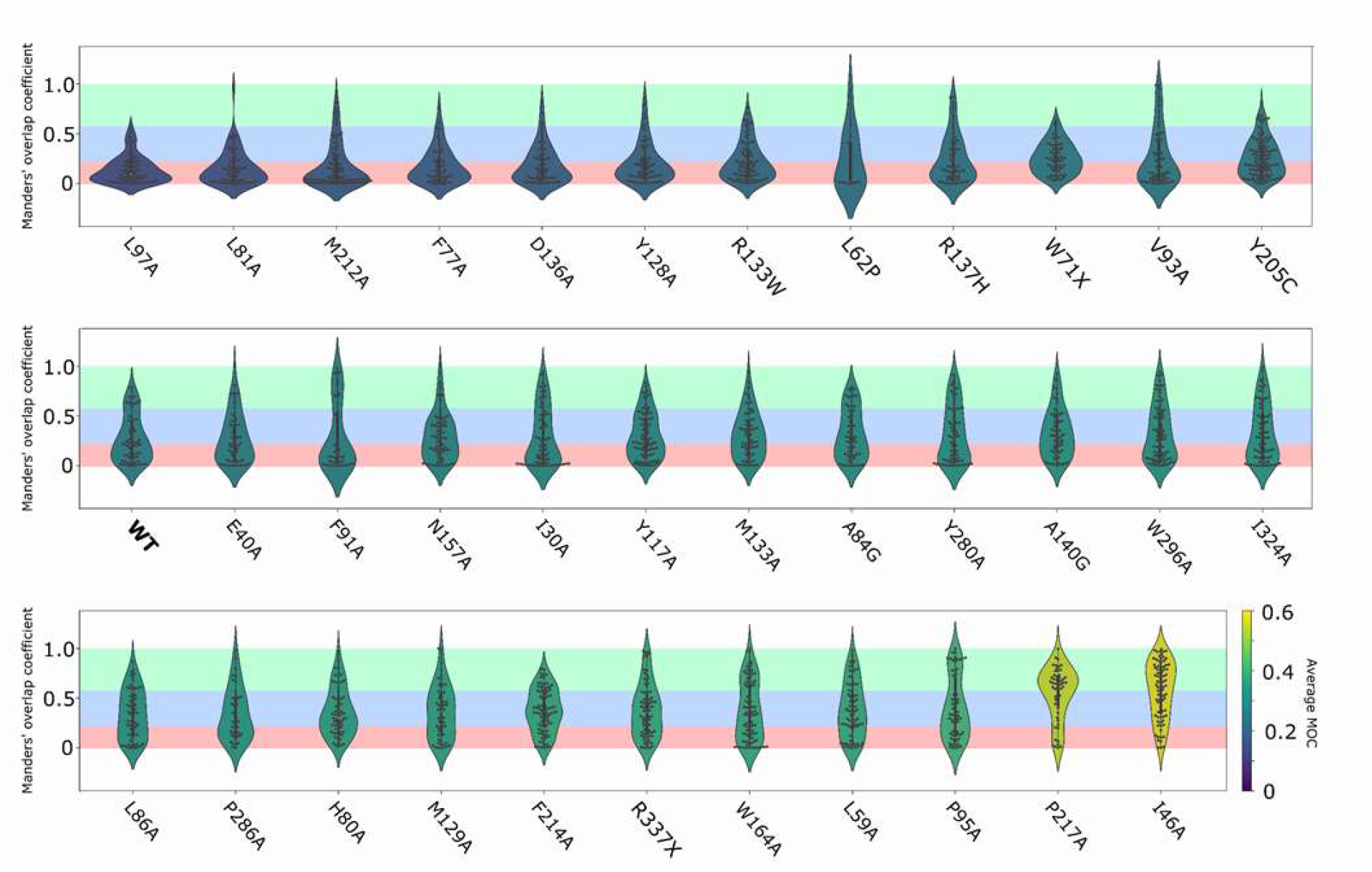
ER localisation per cell for all studied mutants sorted by average ER localisation. Green: high ER localisation, blue: medium ER localisation, red: low ER localisation

**Figure S2:**
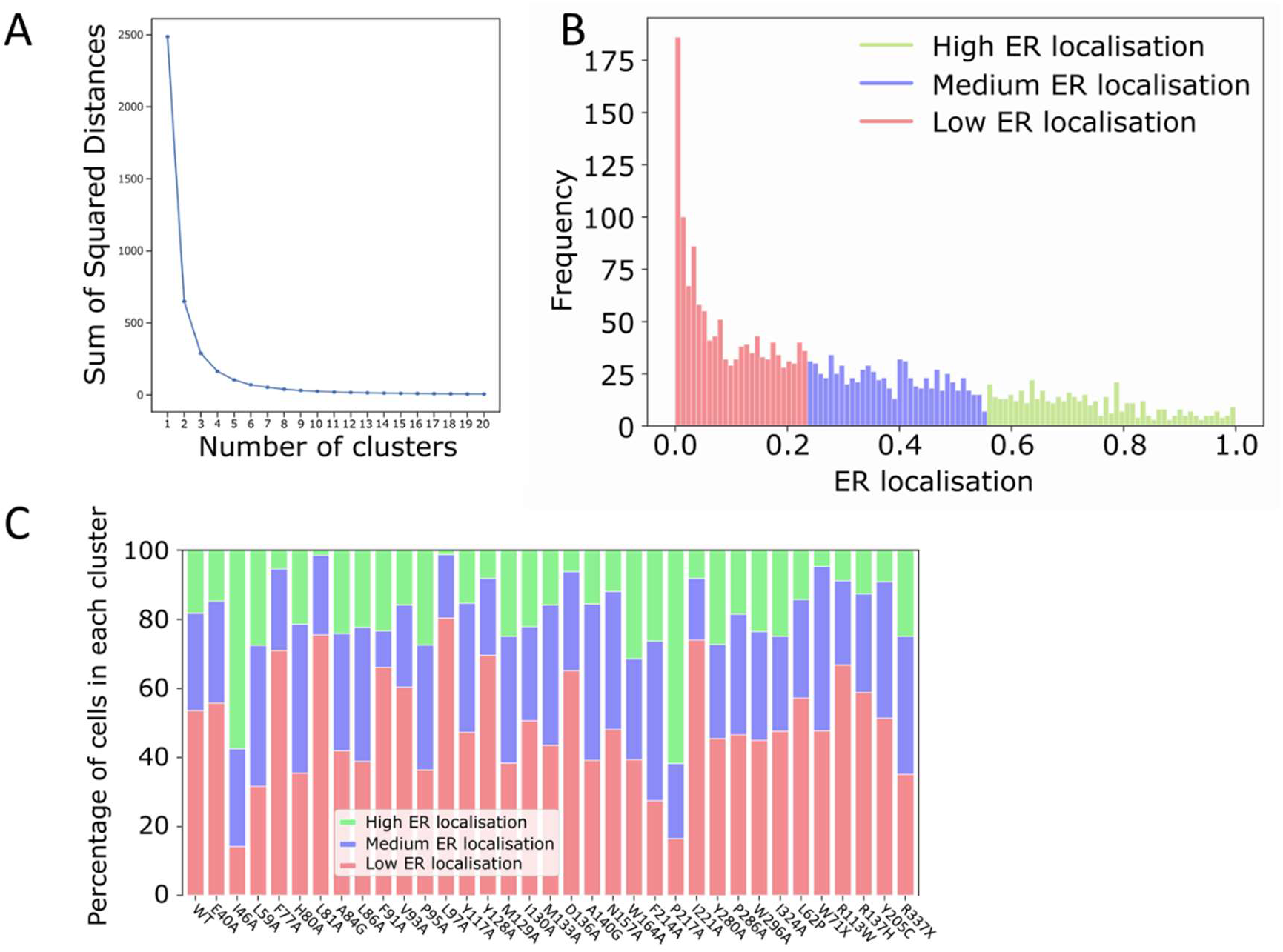
Cells categorised based on ER localisation using k-means clustering A) Elbow plot showing optimum number of clusters B) clusters overlayed with ER localisation distribution of all cells C) percentages of cells in each cluster split out per expressed mutant

**Figure S3:**
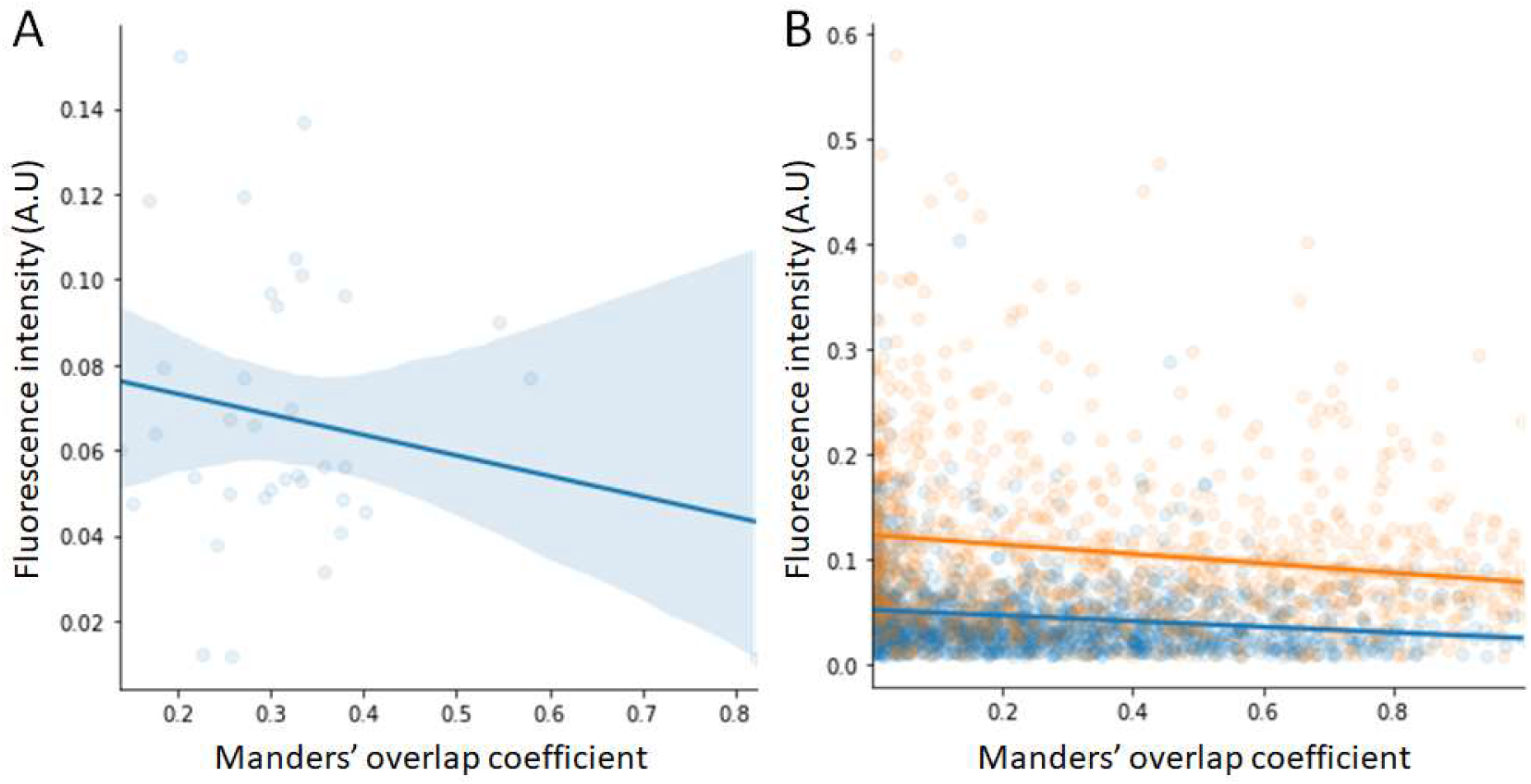
Correlation of V2R Fluorescence Intensity and Manders’ overlap coefficient. A) Average Manders’ overlap coefficient (MOC) and intensity per mutant (R^2 = −4). B) MOC and intensity per cell split out per biological replicate (N=2, R^2 = −0.9 and −1 for red and blue respectively).

**Figure S4:**
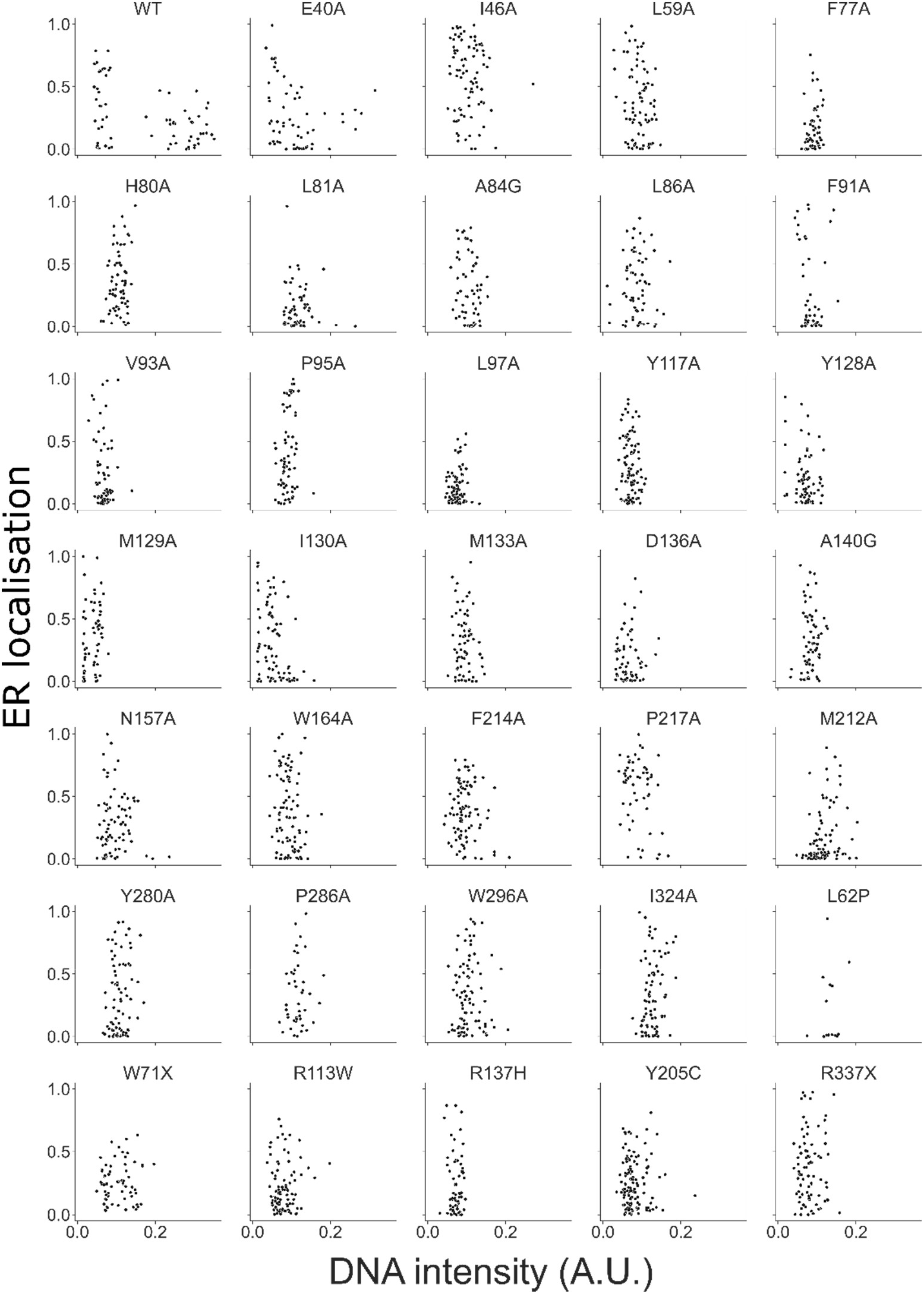
Scatter plots of DAPI fluorescent intensity representing DNA intensity and Manders’ overlap coefficient of V2R with ER markers representing ER localisation of V2R.

